# Structure-based Design of Prefusion-stabilized SARS-CoV-2 Spikes

**DOI:** 10.1101/2020.05.30.125484

**Authors:** Ching-Lin Hsieh, Jory A. Goldsmith, Jeffrey M. Schaub, Andrea M. DiVenere, Hung-Che Kuo, Kamyab Javanmardi, Kevin C. Le, Daniel Wrapp, Alison Gene-Wei Lee, Yutong Liu, Chia-Wei Chou, Patrick O. Byrne, Christy K. Hjorth, Nicole V. Johnson, John Ludes-Meyers, Annalee W. Nguyen, Juyeon Park, Nianshuang Wang, Dzifa Amengor, Jennifer A. Maynard, Ilya J. Finkelstein, Jason S. McLellan

## Abstract

The COVID-19 pandemic caused by the novel coronavirus SARS-CoV-2 has led to accelerated efforts to develop therapeutics, diagnostics, and vaccines to mitigate this public health emergency. A key target of these efforts is the spike (S) protein, a large trimeric class I fusion protein that is metastable and difficult to produce recombinantly in large quantities. Here, we designed and expressed over 100 structure-guided spike variants based upon a previously determined cryo-EM structure of the prefusion SARS-CoV-2 spike. Biochemical, biophysical and structural characterization of these variants identified numerous individual substitutions that increased protein yields and stability. The best variant, HexaPro, has six beneficial proline substitutions leading to ∼10-fold higher expression than its parental construct and is able to withstand heat stress, storage at room temperature, and multiple freeze-thaws. A 3.2 Å-resolution cryo-EM structure of HexaPro confirmed that it retains the prefusion spike conformation. High-yield production of a stabilized prefusion spike protein will accelerate the development of vaccines and serological diagnostics for SARS-CoV-2.

## INTRODUCTION

Coronaviruses are enveloped viruses containing positive-sense RNA genomes. Four human coronaviruses generally cause mild respiratory illness and circulate annually. However, SARS-CoV and MERS-CoV were acquired by humans via zoonotic transmission and caused outbreaks of severe respiratory infections with high case-fatality rates in 2002 and 2012, respectively^1,2^. SARS-CoV-2 is a novel betacoronavirus that emerged in Wuhan, China in December 2019 and is the causative agent of the ongoing COVID-19 pandemic^3,4^. As of May 26, 2020, the WHO has reported over 5 million cases and 350,000 deaths worldwide. Effective vaccines, therapeutic antibodies and small-molecule inhibitors are urgently needed, and the development of these interventions is proceeding rapidly.

Coronavirus virions are decorated with a spike (S) glycoprotein that binds to host-cell receptors and mediates cell entry via fusion of the host and viral membranes^5^. S proteins are trimeric class I fusion proteins that are expressed as a single polypeptide that is subsequently cleaved into S1 and S2 subunits by cellular proteases^6,7^. The S1 subunit contains the receptor-binding domain (RBD), which, in the case of SARS-CoV-2, recognizes the angiotensin-converting enzyme 2 (ACE2) receptor on the host-cell surface^8–10^. The S2 subunit mediates membrane fusion and contains an additional protease cleavage site, referred to as S2′, that is adjacent to a hydrophobic fusion peptide. Binding of the RBD to ACE2 triggers S1 dissociation, allowing for a large rearrangement of S2 as it transitions from a metastable prefusion conformation to a highly stable postfusion conformation^6,11^. During this rearrangement, the fusion peptide is inserted into the host-cell membrane after cleavage at S2′, and two heptad repeats in each protomer associate to form a six-helix bundle that brings together the N- and C-termini of the S2 subunits as well as the viral and host-cell membranes. Attachment and entry are essential for the viral life cycle, making the S protein a primary target of neutralizing antibodies and a critical vaccine antigen^12,13^.

A stabilized prefusion conformation of class I fusion proteins is desirable for vaccine development because this conformation is found on infectious virions and displays most or all of the neutralizing epitopes that can be targeted by antibodies to prevent the entry process^14–16^. We and others have observed that prefusion stabilization tends to increase the recombinant expression of viral glycoproteins, possibly by preventing triggering or misfolding that results from a tendency to adopt the more stable postfusion structure. For example, structure-based design of prefusion-stabilized MERS-CoV and SARS-CoV spike ectodomains resulted in homogeneous preparations of prefusion spikes and greatly increased yields^15^. These variants (S-2P) contained two consecutive proline substitutions in the S2 subunit in a turn between the central helix (CH) and heptad repeat 1 (HR1) that must transition to a single, elongated α-helix in the postfusion conformation. Prefusion-stabilized spike variants are also superior immunogens to wild-type spike ectodomains^15^, and have been used to determine high-resolution spike structures by cryo-EM^17–20^. Importantly, the successful transplantation of this double-proline substitution into the SARS-CoV-2 spike (SARS-CoV-2 S-2P) allowed for the rapid determination of high-resolution cryo-EM structures and accelerated development of vaccine candidates^21,22^. However, even with these substitutions, the SARS-CoV-2 S-2P ectodomain is unstable and difficult to produce reliably in mammalian cells, hampering biochemical research and development of subunit vaccines.

Here, we employ structure-based design to increase the yield and stability of the SARS-CoV-2 spike ectodomain in the prefusion conformation. We report multiple prolines, disulfide bonds, salt bridges, and cavity-filling substitutions that increase expression and/or stability of the spike relative to the S-2P base construct. Combining four proline substitutions into a single construct, termed HexaPro, stabilized the prefusion state and increased expression 10-fold. A high-resolution cryo-EM structure of this variant confirms that the proline substitutions adopt the designed conformations and do not disrupt the conformation of the S2 subunit, thus preserving its antigenicity. This work will facilitate production of prefusion spikes for diagnostic kits and subunit vaccines, and has broad implications for next-generation coronavirus vaccine design.

## RESULTS

### Structure-based design of prefusion-stabilized SARS-CoV-2 spikes

To generate a prefusion-stabilized SARS-CoV-2 spike protein that expresses at higher levels and is more stable than our original S-2P construct^21^ we analyzed the SARS-CoV-2 S-2P cryo-EM structure (PDB ID: 6VSB) and designed substitutions based upon knowledge of class I fusion protein function and general protein stability principles. These strategies included the introduction of disulfide bonds to prevent conformational changes during the pre-to-postfusion transition, salt bridges to neutralize charge imbalances, hydrophobic residues to fill internal cavities, and prolines to cap helices or stabilize loops in the prefusion state. We cloned 100 single S-2P variants and characterized their relative expression levels (**Table S1**), and for those that expressed well we characterized their monodispersity, thermostability, and quaternary structure. Given that the S2 subunit undergoes large-scale refolding during the pre-to-postfusion transition, we exclusively focused our efforts on stabilizing S2. Substitutions of each category were identified that increased expression while maintaining the prefusion conformation (**Fig. 1 and 2A**). Overall, 26 out of the 100 single-substitution variants had higher expression than S-2P (**Table S1**).

**Figure 1.**
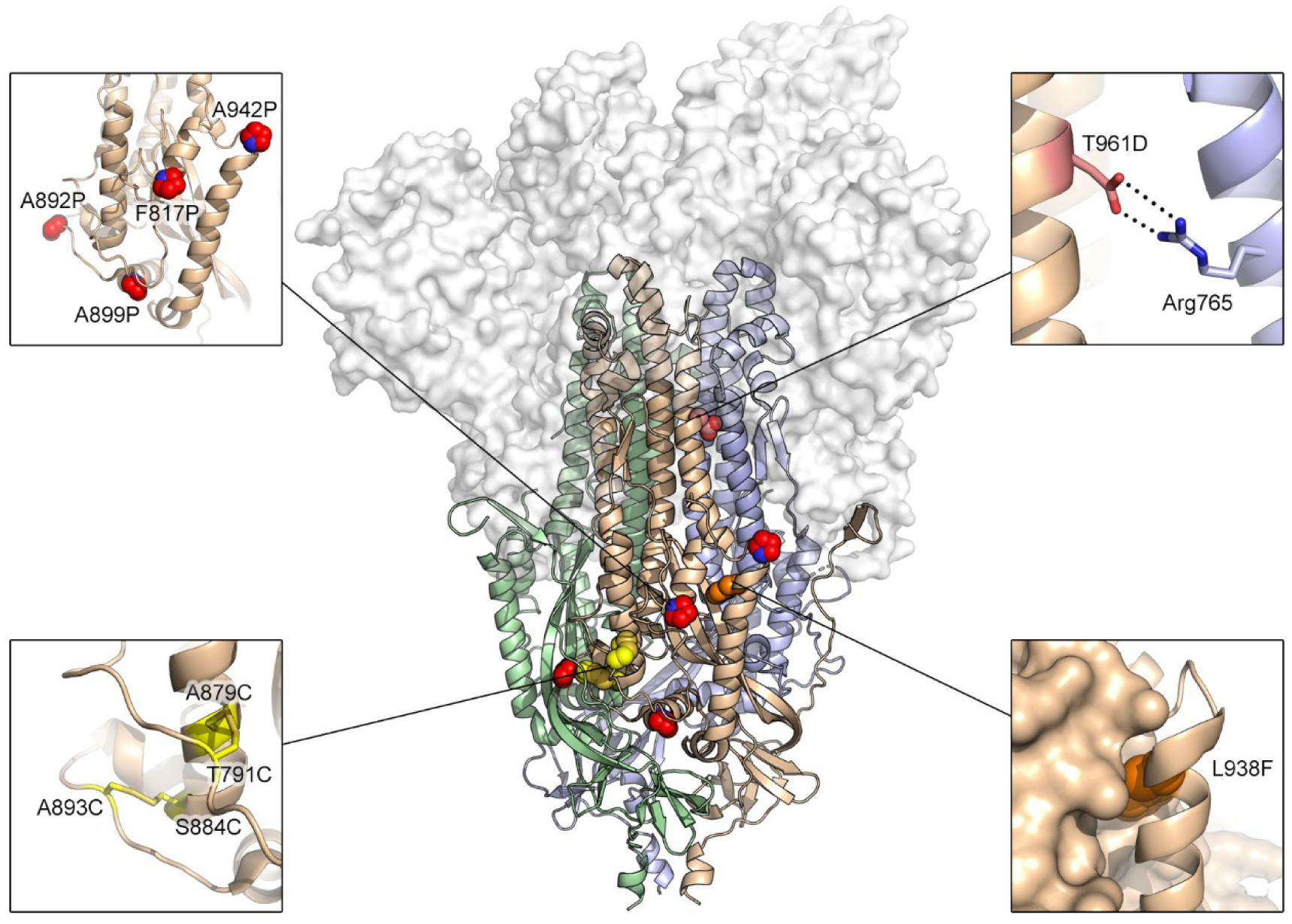
Exemplary substitutions for SARS-CoV-2 spike stabilization. Side view of the trimeric SARS-CoV-2 spike ectodomain in a prefusion conformation (PDB ID: 6VSB). The S1 domains are shown as a transparent molecular surface. The S2 domain for each protomer is shown as a ribbon diagram. Each inset corresponds to one of four types of spike modifications (proline, salt bridge, disulfide, cavity filling). Side chains in each inset are shown as red spheres (proline), yellow sticks (disulfide), red and blue sticks (salt bridge) and orange spheres (cavity filling).

**Figure 2.**
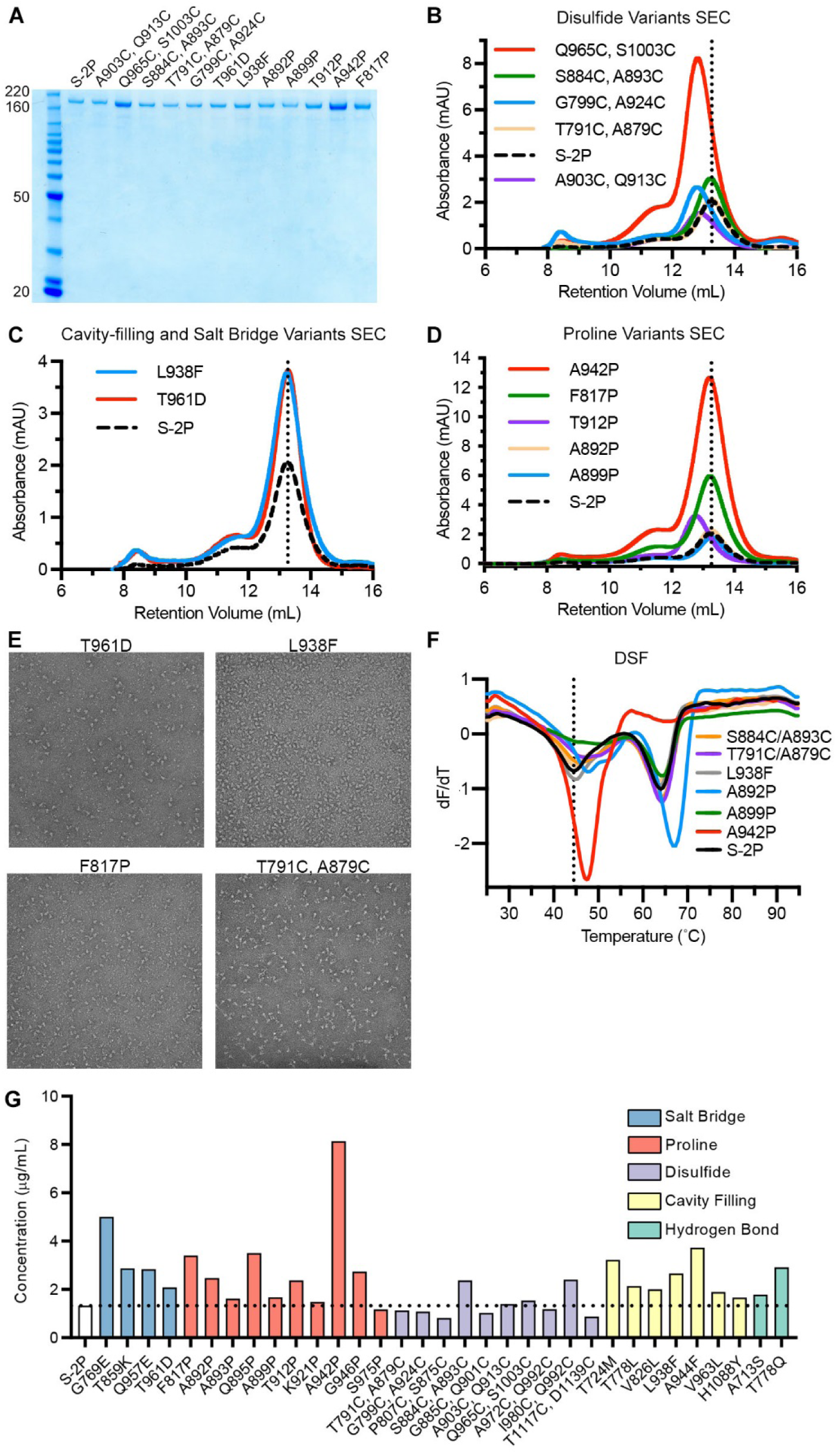
Characterization of single-substitution spike variants. (A) SDS-PAGE of SARS-CoV-2 S-2P and single-substitution spike variants. Molecular weight standards are indicated at the left in kDa. (B-D) Size exclusion chromatography of purified spike variants, grouped by type (B, disulfide variants; C, cavity filling and salt bridge; D, proline). A vertical dotted line indicates the characteristic peak retention volume for S-2P. (E) Representative negative stain electron micrographs for four variants. (F) Differential scanning fluorimetry analysis of spike variant thermostability. The vertical dotted line indicates the first apparent melting temperature for S-2P. (G) Concentrations of individual variants in culture medium, determined by quantitative biolayer interferometry. Variants are colored by type. The horizontal dotted line indicates the calculated concentration of S-2P, which was used as a control for comparison.

### Single-substitution spike variants

One common strategy to stabilize class I fusion proteins, such as the spike, is to covalently link a region that undergoes a conformational change to a region that does not via a disulfide bond. For instance, the Q965C/S1003C substitution attempts to link HR1 to the central helix, whereas G799C/A924C aims to link HR1 to the upstream helix. These two variants boosted protein expression 3.8-fold and 1.3-fold compared to S-2P, respectively (**Fig. 2B**). However, the size-exclusion chromatography (SEC) traces of both variants showed a leftward shift compared to S-2P, indicating that the proteins were running larger than expected, which agreed well with negative stain electron microscopy (nsEM) results that showed partially misfolded spike particles (**Fig. S1**). In contrast, S884C/A893C and T791C/A879C variants eluted on SEC at a volume similar to S-2P and were well-folded trimeric particles by nsEM (**Fig. 2E**). These variants link the same α-helix to two different flexible loops that pack against a neighboring protomer (**Fig. 1**). Notably, S884C/A893C had two-fold higher expression than S-2P with increased thermostability (**Fig. 2F**).

Cavity-filling substitutions and salt bridges can also enhance protein stability without disturbing the overall fold. Cavity filling has been particularly helpful in stabilizing the prefusion conformations of RSV F and HIV-1 Env^23,24^. We found many cavity-filling and salt bridge designs that improved protein expression compared to S-2P (**Fig. 2G**). For example, L938F and T961D both produced ∼2-fold increases in protein yield and maintained the correct quaternary structure (**Fig. 2C and 2E**), although the thermostability of both variants as assessed by differential scanning fluorimetry (DSF) was unchanged compared to S-2P (**Fig. 2F**).

Previous successes using proline substitutions inspired us to investigate 14 individual variants wherein a proline was substituted into flexible loops or the N-termini of helices in the fusion peptide, HR1, and the region connecting them (CR) (**Table 1 and Fig. 1G**). As expected, multiple proline variants boosted the protein expression and increased the thermostability (**Fig. 2D and 2F**). Two of the most successful substitutions, F817P and A942P, exhibited 2.8 and 6.0-fold increases in protein yield relative to S-2P, respectively. The A942P substitution further increased the melting temperature (Tm) by ∼3 °C, and both variants appeared as well-folded trimers by nsEM (**Fig. 2E** and **Fig. S2**).

### Multi-substitution spike variants

We next generated combination (“Combo”) variants that combined the best-performing modifications from our initial screen. The Combo variants containing two disulfide bonds generally expressed 2-fold lower than the single-disulfide variants, suggesting that they interfered with each other (**Table S2**). Adding one disulfide (S884C/A893C) to a single proline variant (F817P) also reduced the expression level, although the quaternary structure of the spikes was well maintained (**Table S2**, Combo40). The beneficial effect of a disulfide bond was most prominent when combined with L938F, a cavity-filling variant. Combo23 (S884C/A893C, L938F) had higher protein yields than either of its parental variants, but the Tm of Combo23 did not increase compared to S884C/A893C (**Fig. S3B**). In addition, mixing one cavity-filling substitution with one proline substitution (Combo20) increased the expression compared to L938F alone (**Table S2**).

Combining multiple proline substitutions resulted in the most drastic increases in expression and stability (**Fig. 3A**). Combo14, containing A892P and A942P, had a 6.2-fold increase in protein yield compared to A892P alone (**Fig. 3B and 3C**). Adding a third proline, A899P, increased thermostability (+1.2 °C Tm) but did not further increase expression (**Fig. 3C**). Combo46 (A892P, A899P, F817P) had a 3.4-fold increase in protein yield and a 3.3 °C rise in Tm as compared to A892P. The most promising variant, Combo47, renamed HexaPro, contains all four beneficial proline substitutions (F817P, A892P, A899P, A942P) as well as the two proline substitutions in S-2P. HexaPro expressed 9.8-fold higher than S-2P, had ∼5 °C increase in Tm, and retained the trimeric prefusion conformation (**Fig. 3D, Fig. S2**). We focused on this construct for additional characterization.

**Figure 3.**
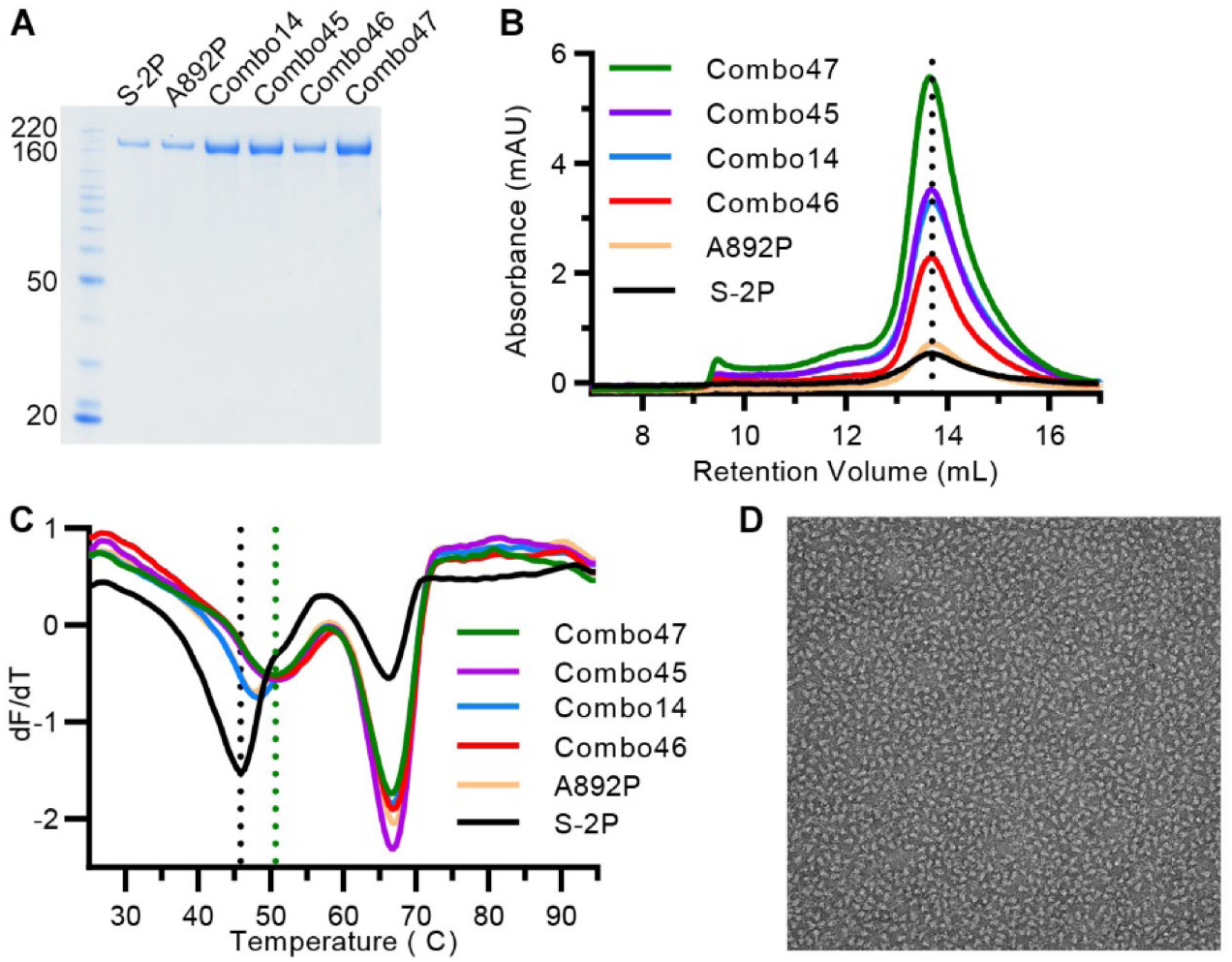
Characterization of multi-substitution spike variants. (A) SDS-PAGE of SARS-CoV-2 Combo variants. Molecular weight standards are indicated at the left in kDa. (B) SEC traces for S-2P, A892P and four Combo variants. The vertical dotted line indicates the peak retention volume for S-2P. (C) DSF analysis of Combo variant thermostability. The black vertical dotted line indicates the first apparent melting temperature for S-2P, the green vertical dotted line shows the first apparent melting temperature for Combo47 (HexaPro). (D) Negative stain electron micrograph of purified Combo47 (HexaPro).

### HexaPro large-scale expression and stress testing

To assess the viability of HexaPro as a potential vaccine antigen or diagnostic reagent, we comprehensively examined large-scale production in FreeStyle 293-F cells, the feasibility of protein expression in ExpiCHO cells, epitope integrity and protein stability. We were able to generate ∼14 mg of HexaPro from 2L of FreeStyle 293-F cells, or 7 mg/L, which represents a greater than 10-fold improvement over S-2P^21^. Importantly, large-scale HexaPro preparations retained a monodisperse SEC peak corresponding to the molecular weight of a glycosylated trimer (**Fig. 4A**) and were indistinguishable from S-2P by nsEM (**Fig. 4B**). Industrial production of recombinant proteins typically relies on CHO cells rather than HEK293 cells. We thus investigated HexaPro expression in ExpiCHO cells via transient transfection. ExpiCHO cells produced 1.3 mg of well-folded protein per 40 mL of culture, or 32.5 mg/L (**Fig. 4C and 4D**). The binding kinetics of HexaPro to the human ACE2 receptor were comparable to that of S-2P (**Fig. 4E and 4F**), with affinities of 13.3 nM and 11.3 nM, respectively. HexaPro remained folded in the prefusion conformation after 3 cycles of freeze-thaw, 2 days incubation at room temperature or 30 minutes at 55 °C (**Fig. 4G and 4H**). In contrast, S-2P showed signs of aggregation after 3 cycles of freeze-thaw, and began unfolding after 30 min at 50 °C. Collectively, these data indicate that HexaPro is a promising candidate for SARS-CoV-2 vaccine development.

**Figure 4.**
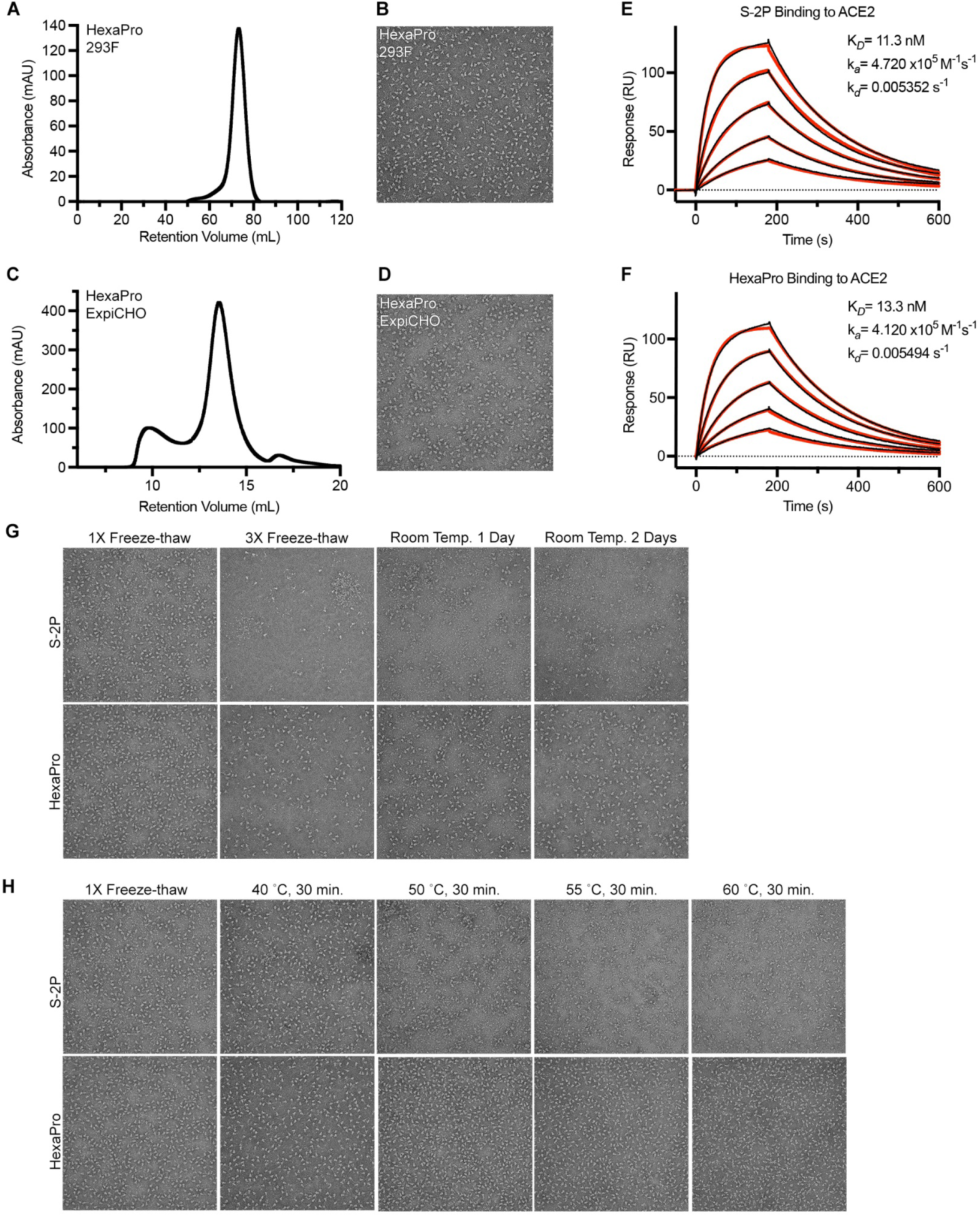
HexaPro exhibits enhanced expression and stability compared to S-2P. (A) SEC trace of a portion of the HexaPro purified from a 2L culture of FreeStyle 293-F cells. (B) Negative stain electron micrograph of HexaPro purified from FreeStyle 293-F cells. (C) SEC trace of HexaPro after purification from a 40 ml culture of ExpiCHO cells. (D) Negative stain electron micrograph of HexaPro purified from ExpiCHO cells. (E-F) Binding of S-2P (E) and HexaPro (F) to human ACE2 assessed by surface plasmon resonance. Binding data are shown as black lines and the best fit to a 1:1 binding model is shown as red lines. (G-H) Assessment of protein stability by negative stain electron microscopy. The top row of micrographs in (G) and (H) corresponds to S-2P, the bottom row corresponds to HexaPro.

### Cryo-EM structure of SARS-CoV-2 S HexaPro

To confirm that our stabilizing substitutions did not lead to any unintended conformational changes, we determined the cryo-EM structure of SARS-CoV-2 S HexaPro. From a single dataset, we were able to obtain high-resolution 3D reconstructions for two distinct conformations of S: one with a single RBD in the up conformation and the other with two RBDs in the up conformation. This two-RBD-up conformation was not observed during previous structural characterization of SARS-CoV-2 S-2P^21,22^. While it is tempting to speculate that the enhanced stability of S2 in HexaPro allowed us to observe this less stable intermediate, validating this hypothesis will require further investigation. Roughly a third (30.6%) of the particles were in the two-RBD-up conformation, leading to a 3.20 Å reconstruction. The remaining particles were captured in the one-RBD-up conformation, although some flexibility in the position of the receptor-accessible RBD prompted us to remove a subset of one-RBD-up particles that lacked clear density for this domain, resulting in a final set of 85,675 particles that led to a 3.21 Å reconstruction (**Fig. 5A, Fig. S4** and **Fig. S5**). Comparison of our one-RBD-up HexaPro structure with the previously determined 3.46 Å S-2P structure revealed an RMSD of 1.2 Å over 436 Cα atoms in S2 (**Fig. 5B**). The relatively high resolution of this reconstruction allowed us to confirm that the stabilizing proline substitutions did not distort the S2 subunit conformation (**Fig. 5C**).

**Figure 5.**
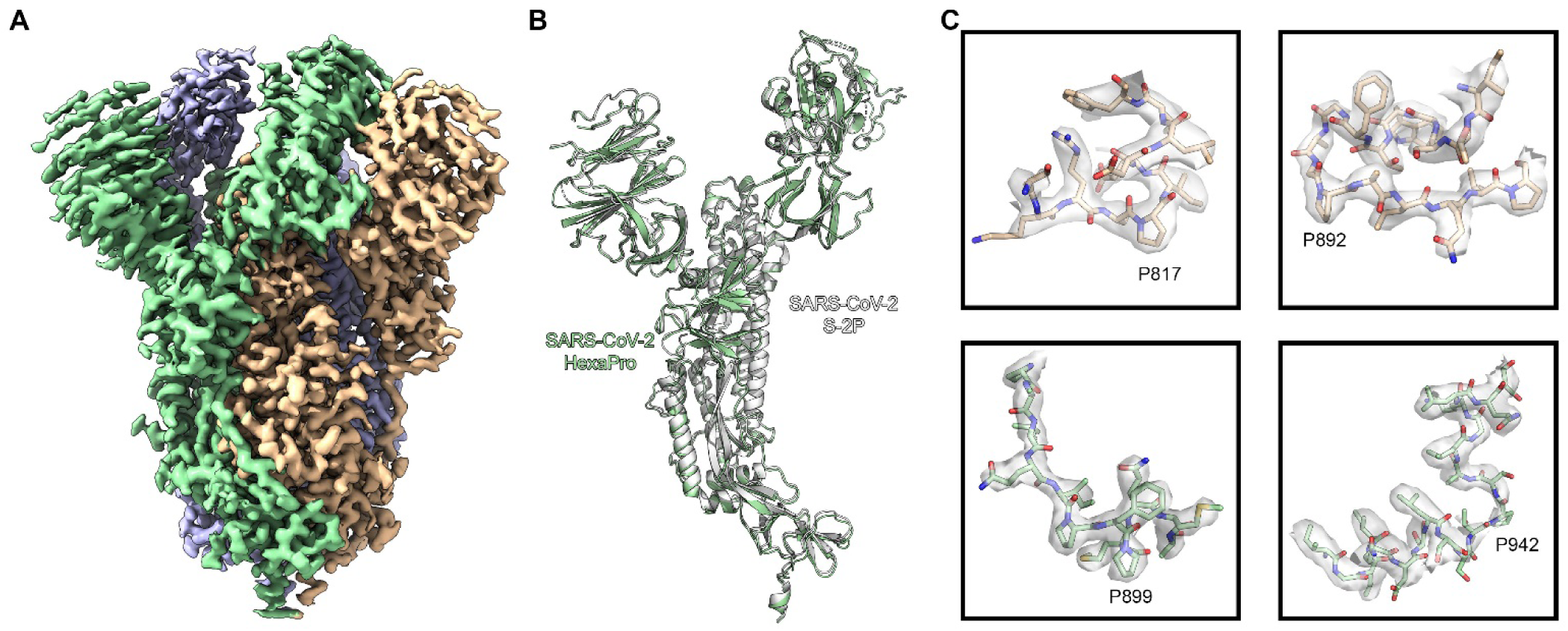
High resolution cryo-EM structure of HexaPro. (A) EM density map of trimeric HexaPro. Each protomer is shown in a different color; the protomer depicted in wheat adopts the RBD-up conformation. (B) Alignment of an RBD-down protomer from HexaPro (green ribbon) with an RBD-down protomer from S-2P (white ribbon, PDB ID: 6VSB). (C) Zoomed view of the four proline substitutions unique to HexaPro. The EM density map is shown as a transparent surface, individual atoms are shown as sticks. Nitrogen atoms are colored blue and oxygen atoms are colored red.

## DISCUSSION

Prefusion-stabilized class I viral fusion proteins generally induce more potent neutralizing antibodies and function as better vaccine antigens than their unstabilized counterparts^14–16^. To respond to the urgent need for preventative countermeasures against the COVID-19 pandemic, we used our prefusion-stabilized SARS-CoV-2 S-2P structure^21^ as a guide to design 100 single substitution variants intended to have increased expression or stability. We focused on engineering the S2 subunit because it undergoes large-scale refolding to facilitate membrane fusion. One of the strategies we employed was the introduction of disulfide bonds wherein at least one cysteine is in a region that changes conformation between the pre- and postfusion states. Although this method has been successful in the case of HIV-1 Env (SOSIP) and RSV F (DS-Cav1)^14,23^, the disulfides we introduced into S2 generally had detrimental effects. For example, inter-subunit disulfides (e.g. S659C/S698C) decreased the protein expression by 60% (**Table S1**), and the Q965C/S1003C substitution led to partially mis-folded spikes (**Fig. 2B**). Inter-protomer disulfides have been shown to improve the trimer integrity of HIV-1 Env and the stability of RSV F^25,26^, but the inter-protomeric T961C/S758C substitution ablated expression (**Table S1**). In contrast, we found that stabilizing the flexible loops located in the protomer interfaces was beneficial. Both S884C/A893C and T791C/A879C increased thermostability or expression and resulted in native trimer structures. It is possible that anchoring flexible loops to a relatively rigid α-helix favors protomer assembly.

Introducing a salt bridge at the HIV-1 gp120–gp41 interface not only boosted the protein expression but also enhanced the binding of trimer-specific antibodies, suggesting improved retention of the native quaternary structure^24^. Based on a similar principle, the T961D substitution was introduced to form an electrostatic interaction with Arg765 from a neighboring protomer (**Fig. 1**). Likewise, the G769E substitution was designed to form an inter-protomeric salt bridge with Arg1014. Both variants increased expression and resembled well-folded trimeric spikes (**Fig. 2E, Fig. S2, Table S1**). In addition to salt bridges, filling loosely packed hydrophobic cores that allow the protein to refold can help stabilize the prefusion state, as shown by previous cavity-filling substitutions in RSV F and HIV-1 Env^14,23,27^. Here, the L938F substitution was designed to fill a cavity formed in part by HR1, the FP and a β-hairpin (**Fig. 1**). This variant had a 2-fold increase in expression (**Fig. 2C**) and appeared to have additive effects when paired with disulfide or proline substitutions (**Table S2)**.

Among the best single-substitution variants we discovered were F817P and A942P (**Fig 2**). By further combining them with A892P and A899P substitutions, we generated the highest expressing construct, HexaPro. This result is reminiscent of previous successful applications of proline substitutions to class I fusion proteins including HIV-1 Env, influenza HA, RSV F, hMPV F, MERS-CoV S, Lassa GPC and Ebola GP^14,15,23,28–31^. Solvent accessibility of hydrophobic residues near the fusion peptide was a concern for influenza HA stem-only designs^32^, and similarly we addressed this issue by replacing the exposed Phe817 with Pro (**Fig. 5C**). The A942P substitution imposes rigidity to the flexible loop between the connector region and HR1, and is similar to that of the T577P substitution found to be helpful for stabilizing Ebola GP^28^.

In our HexaPro cryo-EM dataset we observed a third of the particles in a two-RBD-up conformation. This had not been previously observed for SARS-CoV-2 spikes until a recent structure was determined of a modified spike containing four hydrophobic substitutions that brought subdomain 1 closer to S2^33^. We hypothesize that the more stable S2 in HexaPro allowed us to capture this relatively unstable conformation that may transiently exist prior to triggering and dissociation of S1. This is similar to what was observed in the structures of the MERS-CoV S-2P spike, where even the 3-RBD-up conformation could be observed^15^. Additionally, HexaPro spikes were able to retain the prefusion state after freeze-thaws, room temperature storage, and heat stress, which should aid in the development of HexaPro spikes as subunit vaccine antigens. HexaPro spikes may also improve DNA or mRNA-based vaccines by producing more antigen per nucleic acid molecule, thus improving efficacy at the same dose or maintaining efficacy at lower doses. Finally, we demonstrate that 32 mg of well-folded HexaPro can be obtained from 1L of ExpiCHO cells, indicating a clear path to industrial-level production to meet global demand for this essential SARS-CoV-2 protein.

## ACKNOWLEDGMENTS

We thank members of the Maynard, Finkelstein, and McLellan Laboratories for providing helpful comments on the manuscript. In addition, we would like to thank Dr. Thomas Edwards and Ulrich Baxa for cryo-EM data collection. We also thank Dr. Eric Fich for providing helpful data analysis. This work was supported by NIH grant R01-AI127521 to J.S.M., and grants from the Welch Foundation (F-1808 to I.J.F.), the NIH (GM120554 and GM124141 to I.J.F) and the NSF (1453358 to I.J.F.). I.J.F. is a CPRIT Scholar in Cancer Research. This research was, in part, supported by the National Cancer Institute’s National Cryo-EM Facility at the Frederick National Laboratory for Cancer Research under contract HSSN261200800001E. The Sauer Structural Biology Laboratory is supported by the University of Texas College of Natural Sciences and by award RR160023 from the Cancer Prevention and Research Institute of Texas (CPRIT).

## AUTHOR CONTRIBUTIONS

Conceptualization, C.-L.H. and J.S.M.; Investigation and visualization, C.-L.H., J.A.G., C.-W.C., A.M.D., K.J., H.-C.K., K.C.L., A.G.-W.L., Y.L., J.M.S., D.W., P.O.B., C.K.H., N.V.J., J.L.-M., A.W.N., J.P., and D.A.; Writing – Original Draft, C.-L.H., J.A.G., D.W., P.O.B., C.K.H., N.V.J., and J.S.M; Writing – Reviewing & Editing, C.-L.H., J.A.G., D.W., P.O.B., N.W., C.K.H., N.V.J., J.A.M., I.J.F., and J.S.M.; Supervision, J.A.M., I.J.F. and J.S.M.

## SUPPLEMENTAL FIGURE LEGENDS

## METHODS

### Design scheme for prefusion-stabilized SARS-CoV-2 spike variants

The SARS-CoV-2 S-2P variant was used as the base construct for all subsequent designs^21^. The S-2P base construct comprises residues 1-1208 of SARS-CoV-2 S (GenBank: MN908947) with prolines substituted at residues 986 and 987, “GSAS” substituted at the furin cleavage site (residues 682–685), and C-terminal foldon trimerization motif, HRV3C protease recognition site, Twin-Strep-tag and octa-histidine tag cloned into the mammalian expression plasmid pαH. Using this plasmid as a template, desired mutations were introduced at selected positions within the SARS-CoV-2 S2 subunit. Based on SARS-CoV-2 S-2P cryo-EM structure (PDB ID: 6VSB), pairs of residues with Cβ atoms less than 4.6 Å apart were considered for disulfide bond designs. We particularly targeted the regions that move drastically during the pre- to postfusion transition such as the fusion peptide, connector region and HR1. Salt bridge variants required that the charged groups of the substituted residues were predicted to be within 4.0 Å. For residues in loops, a slightly longer distance than 4.0 Å was allowed. Core-facing residues with sidechains adjacent to a pre-existing internal cavity were examined for potential substitutions to bulkier hydrophobic residues. Proline substitutions were designed in the FP, connector region, or HR1 and placed either in a flexible loop or at the N-terminus of a helix. Combinations were chosen to test whether pairs of the same type of design (e.g. disulfide/disulfide) or different types of designs (e.g. disulfide/proline) could result in additive effects on spike expression and stability.

### Protein expression and purification

Plasmids encoding S variants were transiently transfected into FreeStyle 293-F cells (Thermo Fisher) using polyethyleneimine, with 5 μM kifunensine being added 3h post-transfection. Cultures were harvested four days after transfection and the medium was separated from the cells by centrifugation. Supernatants were passed through a 0.22 µm filter and then over StrepTactin resin (IBA). Spike variants were further purified by size-exclusion chromatography using a Superose 6 10/300 column (GE Healthcare) in a buffer composed of 2 mM Tris pH 8.0, 200 mM NaCl and 0.02% NaN_3_. For initial purification and characterization, single-substitution and combination spike variants were purified from 40 mL cultures. For the 2L HexaPro purification, the size-exclusion column used was a Superose 6 16/600 column (GE Healthcare).

ExpiCHO cells were transiently transfected with a plasmid encoding HexaPro using Expifectamine, and cells were grown for six days at 32 °C according to the manufacturer’s High Titer protocol (Thermo Fisher). Supernatants were then passed through a 0.22 µm filter and batch-purified using IMAC resin (Sigma-Aldrich). The IMAC elution was then purified by size-exclusion chromatography using a Superose 6 10/300 column (GE Healthcare) in a buffer composed of 2 mM Tris pH 8.0, 200 mM NaCl and 0.02% NaN_3_.

### Differential scanning fluorimetry

In a 96-well qPCR plate, solutions were prepared with a final concentration of 5X SYPRO Orange Protein Gel Stain (Thermo Fisher) and 0.25 mg/ml spike. Continuous fluorescence measurements (λ_ex_=465 nm, λ_em_=580 nm) were performed using a Roche LightCycler 480 II, using a temperature ramp rate of 4.4 °C/minute increasing from 22 °C to 95 °C. Data were plotted as the derivative of the melting curve as a function of temperature.

### Negative stain EM

Purified SARS-CoV-2 S variants were diluted to a concentration of 0.04 mg/mL in 2 mM Tris pH 8.0, 200 mM NaCl and 0.02% NaN_3_. Each protein was deposited on a CF-400-CU grid (Electron Microscopy Sciences) that had been plasma cleaned for 30 seconds in a Solarus 950 plasma cleaner (Gatan) with a 4:1 ratio of O_2_/H_2_ and stained using methylamine tungstate (Nanoprobes). Grids were imaged at a magnification of 92,000X (corresponding to a calibrated pixel size of 1.63 Å/pix) in a Talos F200C TEM microscope equipped with a Ceta 16M detector (Thermo Fisher). Stability experiments with S-2P and HexaPro were performed by imaging samples as described above after 3 rounds of snap freezing with liquid nitrogen and thawing, after storing samples at room temperature for 1-2 days, or after incubating at 50 °C, 55 °C, or 60 °C for 30 minutes in a thermal cycler.

### Biolayer interferometry for quantification of protein expression

Plasmids encoding spike variants were transfected into FreeStyle 293-F cells (Thermo Fisher) in 3 mL of medium and harvested four days after transfection. After centrifugation, supernatant was diluted 5-fold with buffer composed of 10 mM HEPES pH 7.5, 150 mM NaCl, 3 mM EDTA, 0.05% Tween 20 and 1 mg/mL bovine serum albumin. Anti-foldon IgG was immobilized to an anti-human Fc (AHC) biosensor (FortéBio) using an Octet RED96e (FortéBio). The IgG loaded biosensor was then dipped into wells containing individual spike variants. A standard curve was determined by measuring 2-fold serial dilutions of purified S-2P at concentrations ranging from 10 μg/mL to 0.16 μg/mL. The data were reference-subtracted, aligned to a baseline after IgG capture and quantified based on a linear fit of the initial slope for each association curve using Octet Data Analysis software v11.1.

### Surface plasmon resonance

His-tagged HexaPro was immobilized to a NiNTA sensorchip (GE Healthcare) to a level of ∼500 response units (RUs) using a Biacore X100 (GE Healthcare) and running buffer composed of 10 mM HEPES pH 8.0, 150 mM NaCl and 0.05% Tween 20. Serial dilutions of purified hACE2 were injected at concentrations ranging from 250 to 15.6 nM. Response curves were fit to a 1:1 binding model using Biacore X100 Evaluation Software (GE Healthcare).

### Cryo-EM sample preparation and data collection

Purified HexaPro was diluted to a concentration of 0.35 mg/mL in 2 mM Tris pH 8.0, 200 mM NaCl, 0.02% NaN_3_ and applied to plasma-cleaned CF-400 1.2/1.3 grids before being blotted for 6 seconds in a Vitrobot Mark IV (Thermo Fisher) and plunge frozen into liquid ethane. 3,511 micrographs were collected from a single grid using a FEI Titan Krios (Thermo Fisher) equipped with a K3 detector (Gatan). Data were collected at a magnification of 81,000x, corresponding to a calibrated pixel size of 1.08 Å/pix. A full description of the data collection parameters can be found in **Table S3**.

### Cryo-EM data processing

Motion correction, CTF-estimation and particle picking were performed in Warp^34^. Particles were then imported into cryoSPARC v2.15.0 for 2D classification, *ab initio* 3D reconstruction, heterogeneous 3D refinement and non-uniform homogeneous refinement^35^. The one-RBD-up reconstruction was subjected to local B-factor sharpening using LocalDeBlur^36^ and the two-RBD-up reconstruction was sharpened in cryoSPARC. Iterative model building and refinement were performed with Coot, Phenix and ISOLDE^37–39^.

---

**Figure S1.**
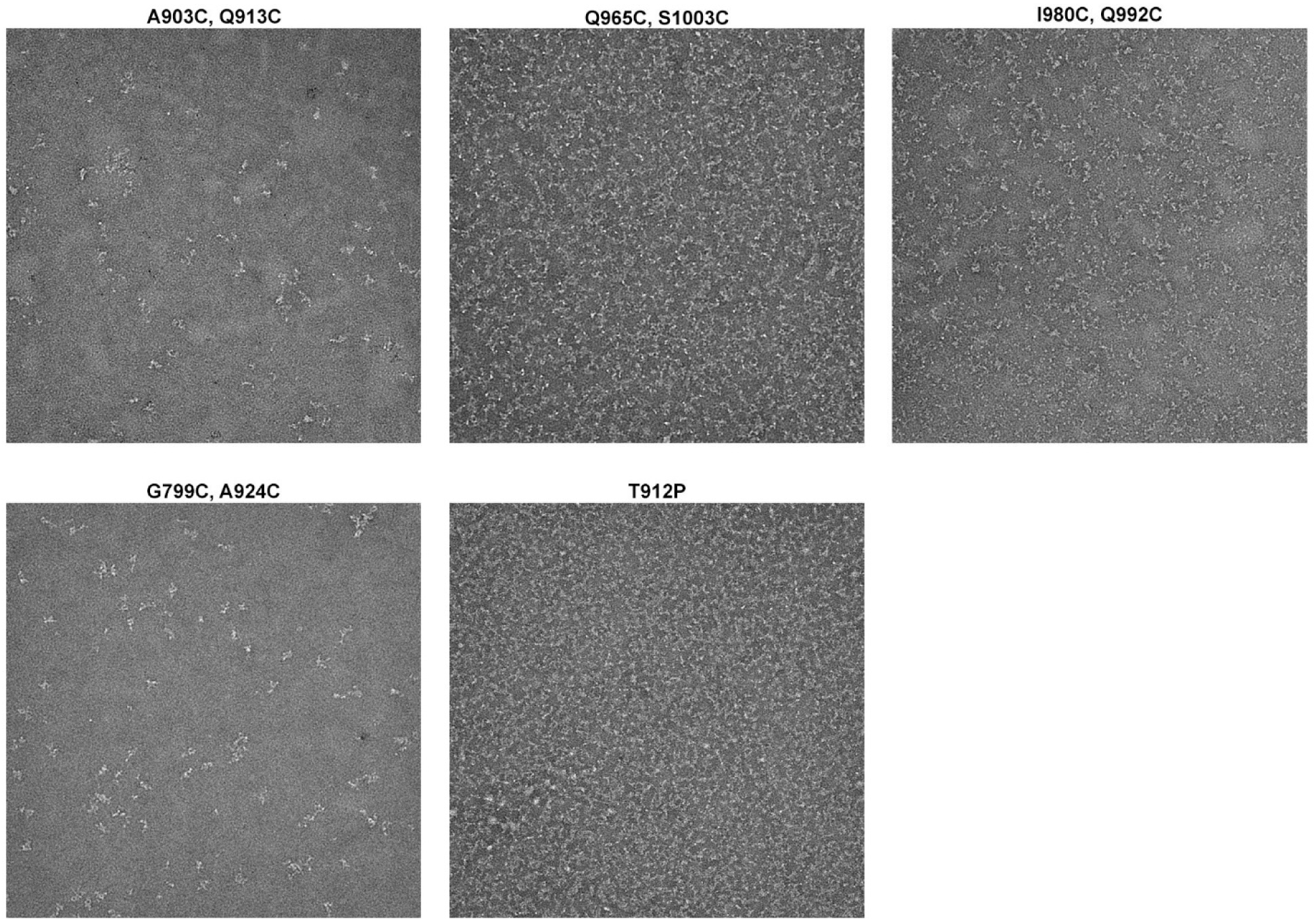
Negative-stain EM images of variants with left-shifted SEC peaks.

**Figure S2.**
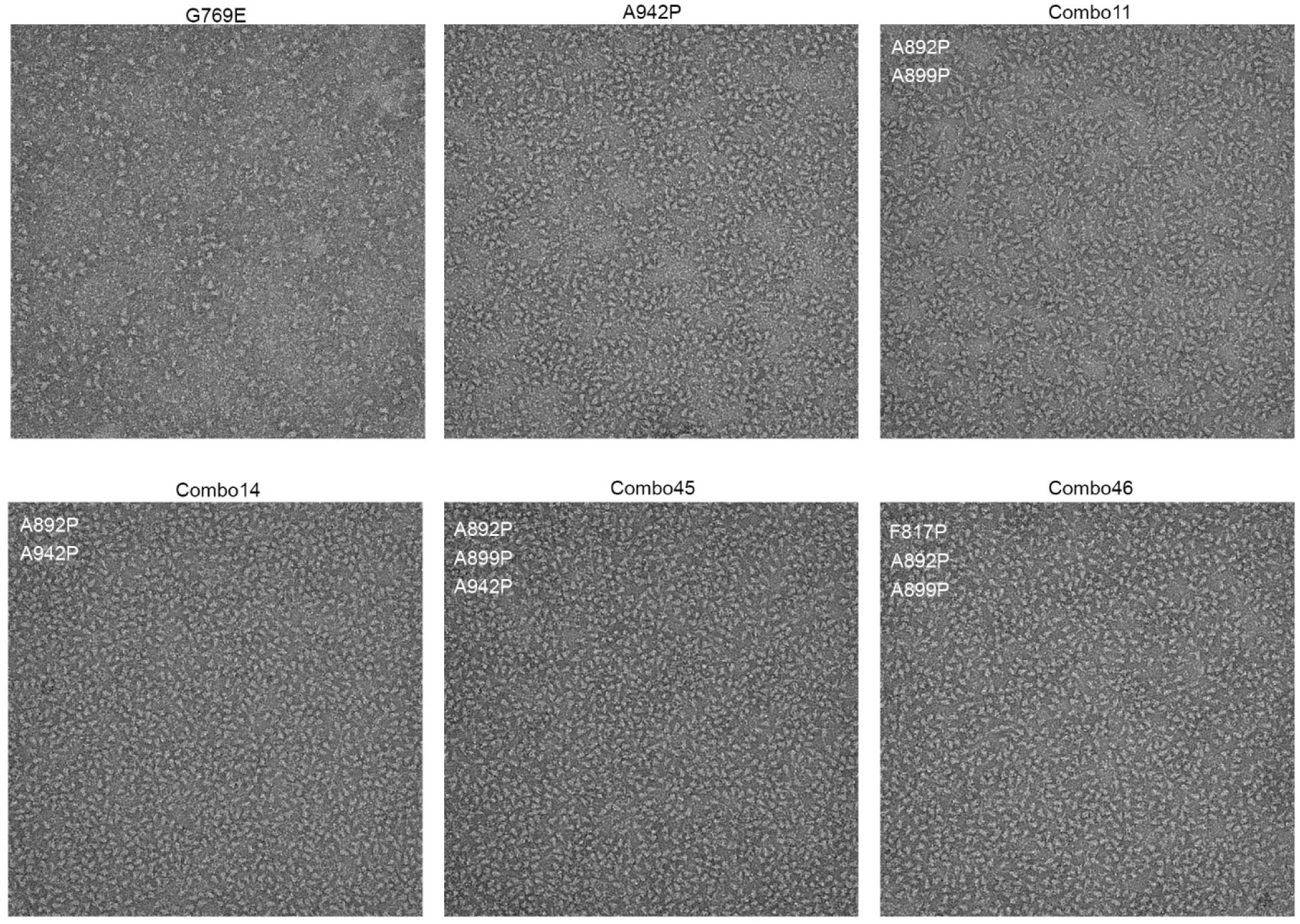
Negative-stain EM images of well-folded particles.

**Figure S3.**
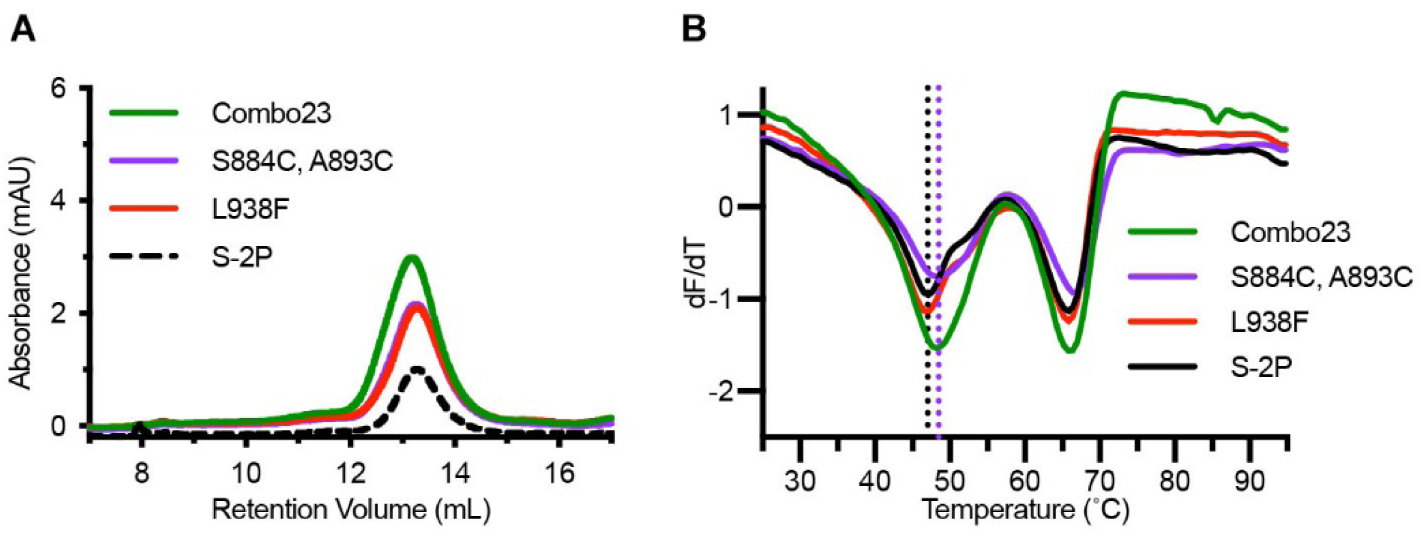
Characterization of a disulfide and cavity-filling combination variant (Combo23). (A) SEC traces of S-2P, Combo23, and the parental variants S884C/A893C (disulfide bond) and L938F (cavity filling). (B) DSF melting temperature analysis of S-2P, Combo23, and its parental variants. The black dashed line represents the Tm of S-2P, and the purple dashed line represents the Tm of S884C/A893C.

**Figure S4.**
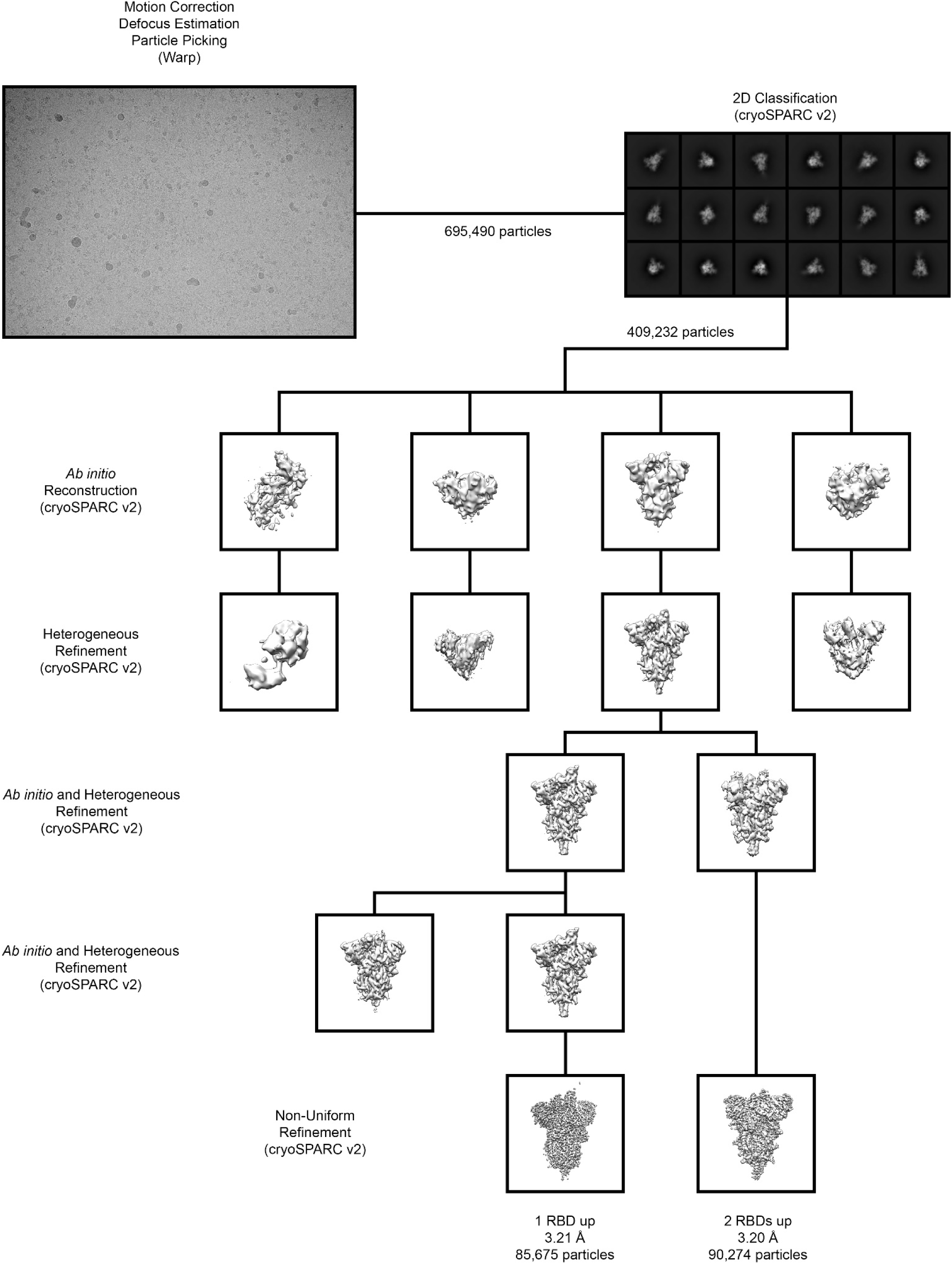
Cryo-EM data processing workflow.

**Figure S5.**
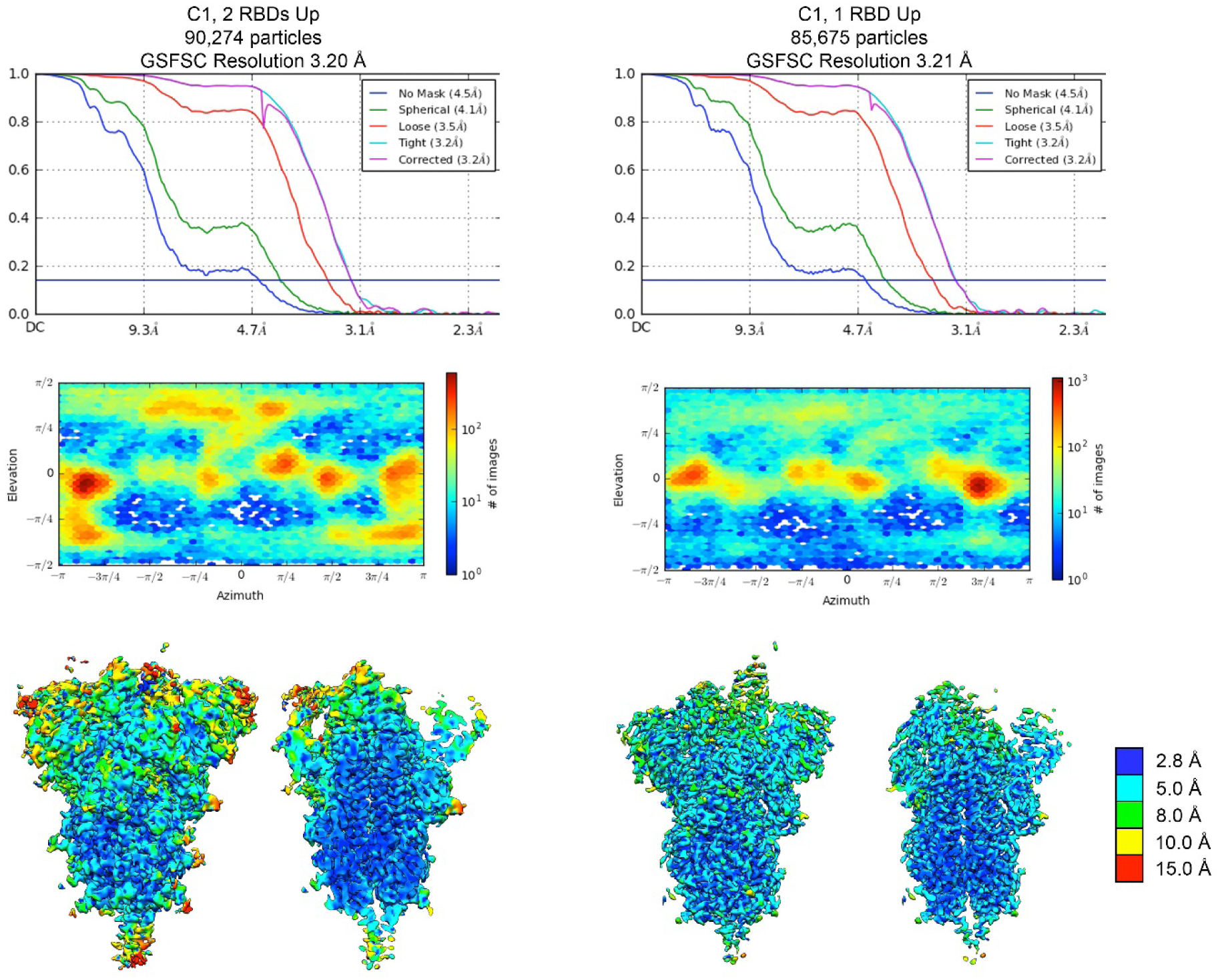
Cryo-EM structure validation. FSC curves and viewing distribution plots, generated in cryoSPARC v2.15, are shown for both the two-RBD-up (*left*) and the one-RBD-up (*right*) reconstruction. Cryo-EM density of each reconstruction is shown and colored according to local resolution, with a central slice through the density shown to the right.

**Table S1.** Expression summary of variants with single substitutions.

| Substitution(s) | Strategy | Fold change in expression relative to S-2P |
| --- | --- | --- |
| T547C, N978C | Disulfide | 0<sup>b</sup> |
| A570C, V963C | Disulfide | 0<sup>b</sup> |
| S659C, S698C | Disulfide | 0.4<sup>a</sup> |
| Replace (673-686) with GS | Remove flexible region | 0<sup>b</sup> |
| Replace (673-686) with GS + A672C, A694C | Disulfide, Remove flexible region | <0.5<sup>b</sup> |
| N703Q, V705C, A893C | Disulfide | <0.5<sup>b</sup> |
| V705C, A893C | Disulfide | <0.5<sup>b</sup> |
| A713S | H bond | 1.0<sup>a</sup> |
| V722C, A930C | Disulfide | <0.1<sup>b</sup> |
| T724M | Cavity-filling | 1.3<sup>a</sup> |
| L727C, S1021C | Disulfide | <0.5<sup>b</sup> |
| P728C, V951C | Disulfide | 0<sup>b</sup> |
| V729C, A1022C | Disulfide | <0.1<sup>a</sup> |
| S730L | Cavity-filling | 0<sup>b</sup> |
| S730R | Salt bridge | 0.15<sup>a</sup> |
| S735C, T859C | Disulfide | <0.5<sup>b</sup> |
| V736C, L858C | Disulfide | 0<sup>b</sup> |
| T752K | Salt bridge | <0.5<sup>b</sup> |
| A766E | Salt bridge | <0.5<sup>b</sup> |
| G769E | Salt bridge | 3.0<sup>a</sup> |
| I770C, A1015C | Disulfide | <0.5<sup>b</sup> |
| T778Q | Hydrogen bond | 2.6<sup>a</sup> |
| T778L | Cavity-filling | 1.5<sup>a</sup> |
| T791C, A879C | Disulfide | 1.0<sup>b</sup> |
| G799C, A924C | Disulfide | 1.3<sup>a</sup> |
| P807C, S875C | Disulfide | 1.1<sup>a</sup> |
| F817P | Proline | 2.8<sup>a</sup> |
| E819C, S1055C | Disulfide | 0<sup>b</sup> |
| E819C, Q1054C | Disulfide | 0<sup>b</sup> |
| L822C, A1056C | Disulfide | 0<sup>b</sup> |
| V826L | Cavity-filling | 1.0<sup>b</sup> |
| L828K | Salt bridge | 0.8<sup>a</sup> |
| L828R | Salt bridge | 0.4<sup>a</sup> |
| Δ(829-851) | Remove flexible region | <0.5<sup>b</sup> |
| T859K | Salt bridge | 2.1<sup>a</sup> |
| P862E | Salt bridge | <0.5<sup>b</sup> |
| L865P, Q779M | Proline, cavity-filling | <0.5<sup>b</sup> |
| T866P | Proline | <0.5<sup>b</sup> |
| I870C, S1055C | Disulfide | 0<sup>b</sup> |
| T874C, S1055C | Disulfide | <0.5<sup>b</sup> |
| S875F | Cavity-filling | <0.5<sup>b</sup> |
| S884C, A893C | Disulfide | 1.5<sup>a</sup> |
| G885C, Q901C | Disulfide | 1.1<sup>a</sup> |
| G889C, L1034C | Disulfide | <0.1<sup>a</sup> |
| A890V | Cavity-filling | 1.0<sup>b</sup> |
| A892P | Proline, cavity-filling | 1.0<sup>a</sup> |
| A893P | Proline | 1.5<sup>b</sup> |
| L894F | Cavity-filling | 0.9<sup>a</sup> |
| Q895P | Proline | 2.1<sup>b</sup> |
| I896C, Q901C | Disulfide | 0<sup>b</sup> |
| A899F | Cavity-filling | 0.3<sup>b</sup> |
| A899P | Proline, Cav | 0.84<sup>a</sup> |
| Q901M | Cavity-filling | 0.9<sup>a</sup> |
| A903C, Q913C | Disulfide | 0.82<sup>b</sup> |
| V911C, N1108C | Disulfide | 0<sup>b</sup> |
| T912R | Salt bridge | <0.5<sup>b</sup> |
| T912P | Proline cavity-filling | 1.5<sup>a</sup> |
| K921P | Proline | 1.1<sup>b</sup> |
| L922P | Proline | 0.8<sup>b</sup> |
| L938F | Cavity-filling | 2.0<sup>a</sup> |
| A942P | Proline | 6.0<sup>a</sup> |
| A944F | Cavity-filling | 1.0<sup>a</sup> |
| A944F, T724I | Cavity-filling | 0.4<sup>a</sup> |
| A944Y | Cavity-filling | 1.9<sup>b</sup> |
| G946P | Proline | 1.0<sup>b</sup> |
| Q957E | Salt bridge | 1.0<sup>a</sup> |
| T961D | Salt bridge | 1.8<sup>a</sup> |
| T961C, S758C | Disulfide | 0<sup>b</sup> |
| T961C, Q762C | Disulfide | 0<sup>b</sup> |
| V963L | Cavity-filling | 1.8<sup>a</sup> |
| Q965C, S1003C | Disulfide | 3.8<sup>a</sup> |
| A972C, Q992C | Disulfide | 1<sup>a</sup> |
| A972C, I980C | Disulfide | 1.3<sup>a</sup> |
| S974C, D979C | Disulfide | 0.3<sup>b</sup> |
| S975P | Proline | 2.2<sup>b</sup> |
| N978P | Proline | 0.9<sup>b</sup> |
| I980C, Q992C | Disulfide | 2.0<sup>a</sup> |
| R1000Y | Cavity-filling + hydrogen bond | 0.3<sup>a</sup> |
| R1000W | Cavity-filling | 1.0<sup>a</sup> |
| S1003V | Cavity-filling | 1.9<sup>b</sup> |
| I1013F | Cavity-filling | 0.8<sup>a</sup> |
| R1039F | Charge removal, pi-pi stacking | 0.5<sup>b</sup> |
| V1040F | Cavity-filling | <0.5<sup>b</sup> |
| V1040Y | Cavity-filling | 0.3<sup>a</sup> |
| H1058W | Cavity-filling | <0.5<sup>b</sup> |
| H1058F | Cavity-filling | 0<sup>b</sup> |
| H1058Y | Cavity-filling | 0.3<sup>a</sup> |
| A1078C, V1133C | Disulfide | <0.5<sup>b</sup> |
| A1080C, I1132C | Disulfide | <0.5<sup>b</sup> |
| I1081C, N1135C | Disulfide | 0.3<sup>a</sup> |
| H1088Y | Cavity-filling | 1.6<sup>a</sup> |
| H1088W | Cavity-filling | 0.6<sup>a</sup> |
| F1103C, P1112C | Disulfide | 0.15<sup>a</sup> |
| V1104I | Cavity-filling | 0.7<sup>a</sup> |
| T1116C, Y1138C | Disulfide | 0<sup>b</sup> |
| T1117C, D1139C | Disulfide | 1.0<sup>a</sup> |
| D1118F | Charge removal, pi-pi stacking | 0.5<sup>b</sup> |
| I1130Y | Hydrogen bond | 0<sup>b</sup> |
| L1141F | Cavity-filling | 0.8<sup>a</sup> |
| ΔHR2 (Δ1161-1208) | Remove flexible region | 2.5<sup>a</sup> |

<sup>a</sup>Quantified using the area under the curve of the size-exclusion trimer peak
<sup>b</sup>Quantified using SDS-PAGE band intensity

**Table S2.** Expression summary of Combo variants.

| Combo # | Substitutions | Strategy | Fold change in expression relative to S-2P |
| --- | --- | --- | --- |
| Combo1 | A903C, Q913C, Q965C, S1003C | Disulfide+Disulfide | 2.2 |
| Combo2 | S884C, A893C, A903C, Q913C | Disulfide+Disulfide | 0.8 |
| Combo3 | T791C, A879C, A903C, Q913C | Disulfide+Disulfide | 0.5 |
| Combo4 | G799C, A924C, A903C, Q913C | Disulfide+Disulfide | 0.5 |
| Combo8 | T791C, A879C, S884C, A893C | Disulfide+Disulfide | 0.5 |
| Combo9 | G799C, A924C, S884C, A893C | Disulfide+Disulfide | 0.4 |
| Combo11 | A892P, A899P | Proline+Proline | 1.9 |
| Combo12 | A892P, T912P | Proline+Proline | 2.7 |
| Combo14 | A892P, A942P | Proline+Proline | 6.2 |
| Combo16 | A899P, A942P | Proline+Proline | 5.1 |
| Combo19 | L938F, A892P | Cavity-filling+Proline | 3.0 |
| Combo20 | L938F, A899P | Cavity-filling+Proline | 3.0 |
| Combo21 | F817P, L938F | Proline+Proline | 3.9 |
| Combo22 | L938F, A942P | Cavity-filling+Proline | 6.0 |
| Combo23 | S884C, A893C, L938F | Disulfide+Cavity-filling | 2.9 |
| Combo24 | T791C, A879C, L938F | Disulfide+Cavity-filling | 2.2 |
| Combo26 | L938F, A903C, Q913C | Cavity-filling+Disulfide | 2.0 |
| Combo40 | F817P, S884C, A893C | Proline+Disulfide | 2.0 |
| Combo42 | T791C, A879C, F817P | Disulfide+Proline | 1.4 |
| Combo45 | A892P, A899P, A942P | 3X Proline | 6.2 |
| Combo46 | F817P, A892P, A899P | 3X Proline | 3.8 |
| Combo47 | F817P, A892P, A899P, A942P | 4X Proline | 9.8 |

**Table S3.** Cryo-EM data collection and refinement statistics.

| **EM data collection and reconstruction statistics** |  |  |
| --- | --- | --- |
| Protein | SARS-CoV-2 S HexaPro | SARS-CoV-2 S HexaPro |
|  | One RBD Up | Two RBDs Up |
| EMDB |  |  |
| Microscope | FEI Titan Krios | FEI Titan Krios |
| Voltage (kV) | 300 | 300 |
| Detector | Gatan K3 | Gatan K3 |
| Magnification | 81,000 | 81,000 |
| Pixel size (Å/pix) | 1.08 | 1.08 |
| Frames per exposure | 40 | 40 |
| Exposure (e<sup>-</sup>/Å<sup>2</sup>) | 45 | 45 |
| Defocus range (μm) | 0.8-2.3 | 0.8-2.3 |
| Micrographs collected | 2,436 | 2,436 |
| Particles extracted/final | 695,490 / 85,675 | 695,490 / 90,274 |
| Symmetry imposed | n/a (C1) | n/a (C1) |
| Masked resolution at 0.143 FSC (Å) | 3.21 | 3.20 |
| **Model refinement and validation statistics** |  |  |
| PDB |  |  |
| Composition |  |  |
| Amino acids | 2,920 |  |
| Glycans | 50 |  |
| RMSD bonds (Å) | 0.007 |  |
| RMSD angles (°) | 0.995 |  |
| Mean B-factors |  |  |
| Amino acids | 37.8 |  |
| Glycans | 55 |  |
| Ramachandran |  |  |
| Favored (%) | 95.4 |  |
| Allowed (%) | 4.5 |  |
| Outliers (%) | 0.1 |  |
| Rotamer outliers (%) | 3.7 |  |
| Clash score | 6.72 |  |
| C-beta outliers (%) | 0.0 |  |
| CaBLAM outliers (%) | 2.75 |  |
| MolProbity score | 2.12 |  |
| EMRinger score | 2.87 |  |

